# Pathway-Specific Biomarker Alterations Decades After Preeclampsia: Associations With Long-Term Cardiorenal Risk

**DOI:** 10.64898/2026.11.17.767451

**Authors:** Vladimir Dokic, Jennet Hatamova, Lisa Vaughan, Muthuvel Jayachandran, Andrea Kattah, Jose Arriola-Montenegro, Elizabeth Ann L. Enninga, Oscar Garcia Valencia, Navin Gupta, Sonja Suvakov, Vesna D. Garovic

**Affiliations:** Division of Nephrology & Hypertension, Mayo Clinic, Rochester, MN USA; Department of Quantitative Health Sciences, Mayo Clinic, Rochester, MN USA; Department of Physiology and Biomedical Engineering, Mayo Clinic College of Medicine and Science, Rochester, MN USA; Division of Hematology, Mayo Clinic College of Medicine and Science, Rochester, MN USA; Department of Obstetrics and Gynecology, Mayo Clinic, Rochester, MN USA

**Keywords:** Preeclampsia, cardiovascular disease, kidney disease, biomarkers

## Abstract

**Background:** Preeclampsia, a heterogeneous disorder involving multiple pathophysiologic pathways, is associated with increased cardiovascular and kidney disease later in life. Whether abnormalities in distinct biological pathways implicated during preeclamptic pregnancies persist decades after delivery, and whether they are differentially associated with subsequent cardiorenal outcomes, remains unknown. We evaluated late-life biomarkers reflecting renal tubular integrity, cellular stress, nitric oxide metabolism, and biological aging in women with and without a history of preeclampsia.

**Methods and Results:** We studied 80 postmenopausal women with a history of preeclampsia (n=40) or normotensive pregnancy (n=40), matched by age and parity. Plasma uromodulin (UMOD), serum growth differentiation factor-15 (GDF-15), serum asymmetric dimethylarginine (ADMA), and leukocyte telomere length were measured in samples collected in 2011–2013. Compared with controls, women with prior preeclampsia had lower UMOD (124±62 versus 167±59 ng/mL; P=0.003), higher log2-transformed ADMA (6.83±0.52 versus 6.49±0.50; P=0.005), and shorter telomere length (6.62±1.68 versus 7.67±1.62 kb; P=0.006); these associations persisted after adjustments for body mass index and anti-hypertensive medication use. GDF-15 was higher in unadjusted analysis but not after adjustment. During a median follow-up of 11.2 years, 32 incident outcomes (coronary artery disease, heart failure, chronic kidney disease, transient ischemic attack, and stroke) occurred. Prior preeclampsia was associated with greater risk of the composite outcome (hazard ratio [HR], 2.43 [95% CI, 1.17–5.06]). Higher GDF-15 was associated with greater risk of the composite outcome after adjustments for body mass index and anti-hypertensive medication use (HR per doubling, 2.08 [95% CI, 1.31–3.28]).

**Conclusions:** A history of preeclampsia was associated decades later with differences in biomarkers reflecting renal tubular integrity, nitric oxide metabolism, and biological aging, as well as increased subsequent cardiorenal risk. GDF-15 was associated with incident cardiovascular or kidney disease and may identify a late-life cellular-stress phenotype associated with increased cardiorenal risk after preeclampsia.

## INTRODUCTION

Preeclampsia (PE) is a heterogeneous syndrome in which multiple, potentially overlapping pathophysiologic pathways contribute to its clinical manifestations.^1^ Abnormalities involving renal function, cellular stress, endothelial and nitric oxide biology, and cellular aging have been described during or around pregnancies complicated by PE.^2–6^ However, it remains uncertain whether abnormalities representing these distinct biological pathways are resolved after delivery or remain detectable decades later. Moreover, whether persistence of abnormalities in different pathways is associated with subsequent cardiovascular and kidney outcomes has not been established.

We therefore selected biomarkers representing distinct biological pathways previously implicated in PE and relevant to long-term cardiorenal health. Circulating uromodulin (UMOD), produced exclusively by renal tubular epithelial cells, reflects functional tubular mass and renal tubular integrity and has been associated with kidney and cardiovascular outcomes.^2,3^ Growth differentiation factor-15 (GDF-15) is a stress-responsive cytokine associated with cellular stress, metabolic stress, senescence, aging, and adverse cardiovascular outcomes.^4^ Asymmetric dimethylarginine (ADMA) is an endogenous inhibitor of nitric oxide synthase and provides a marker of impaired nitric oxide bioavailability and endothelial dysfunction.^5^ Leukocyte telomere length provides a complementary measure of cellular replicative history and biological aging.^6^ Importantly, these pathways have been implicated in the pathophysiology of PE during or around the affected pregnancy, providing a rationale for evaluating whether corresponding abnormalities remain detectable later in life.

We compared plasma UMOD, serum GDF-15, serum ADMA, and leukocyte telomere length in postmenopausal women with a documented history of PE and age- and parity-matched women with normotensive pregnancies. We hypothesized that abnormalities across multiple biological pathways implicated in PE would remain detectable decades after the affected pregnancy, although their relationships with long-term clinical outcomes might differ. We first examined whether prior PE was associated with a late-life biomarker profile characterized by lower UMOD, higher GDF-15 and ADMA, and shorter telomere length. In secondary analyses, we evaluated the relationships of these biomarkers with coronary artery calcification, systemic inflammation, and insulin resistance. Finally, using extended longitudinal follow-up, we examined whether a PE history and biomarkers representing these distinct biological pathways were differentially associated with subsequent cardiovascular and kidney events.

## MATERIALS AND METHODS

### Study Design and Participants

This study was conducted with approval from the Institutional Review Board at Mayo Clinic (Rochester, MN USA). Women who delivered between 1/1/1976 and 12/31/1982 were identified through the Rochester Epidemiology Project medical records-linkage system, as previously described.^7–9^ Participants were matched by age and parity and included women with histories of PE (PE group, n = 40) and women with normotensive pregnancies (control group, n = 40). Medical records were reviewed to apply exclusion criteria, which included a prior history of cardiovascular disease, including coronary artery disease (CAD) and heart failure (HF), stroke, dementia, malignancy (except nonmelanoma skin cancer), autoimmune disorders (e.g., systemic lupus erythematosus), or neurological diseases (e.g., multiple sclerosis). A flowchart of inclusion and exclusion criteria, allocation to the groups and timeline of the experiments is shown in **Figure 1**. Demographic and clinical characteristics of these study subjects at baseline have been reported elsewhere and are shown (with some additions) in **Supplementary Table S1**.^10–12^ A history of PE was confirmed in participants who experienced at least one affected pregnancy during 1976–1982 based on established diagnostic criteria, including: (1) two or more systolic blood pressure (SBP) measurements > 140 mm Hg and/or a diastolic blood pressure (DBP) >90 mm Hg obtained at least 4 hours apart after 20 weeks of gestation; and (2) new-onset proteinuria, defined as a urine dipstick reading ≥ 1+, proteinuria excretion ≥300 mg in a 24-hour urine collection, or a protein-to-creatinine ratio equivalent to ≥ 300 mg per 24 hours.^13^ A PE diagnosis was reconstructed from contemporaneous records by investigators blinded to patient biomarker/outcome status. The study combined a cross-sectional biomarker analysis at the 2011–2013 study visit with prospective ascertainment of incident cardiovascular and kidney outcomes after that visit in the following years. We specifically collected information via chart review regarding CAD, HF, chronic kidney disease (CKD), transient ischemic attack (TIA), and stroke (e.g., the composite event in analyses that followed) that occurred between the time of a woman’s enrollment in the study until her last follow-up date (the time of a woman’s last visit to a Rochester Epidemiology Project-affiliated provider) or death, whichever came first. Follow-up was extended beyond our previous publication on this cohort from 2021^10^ through March 2026.

**Figure 1.**
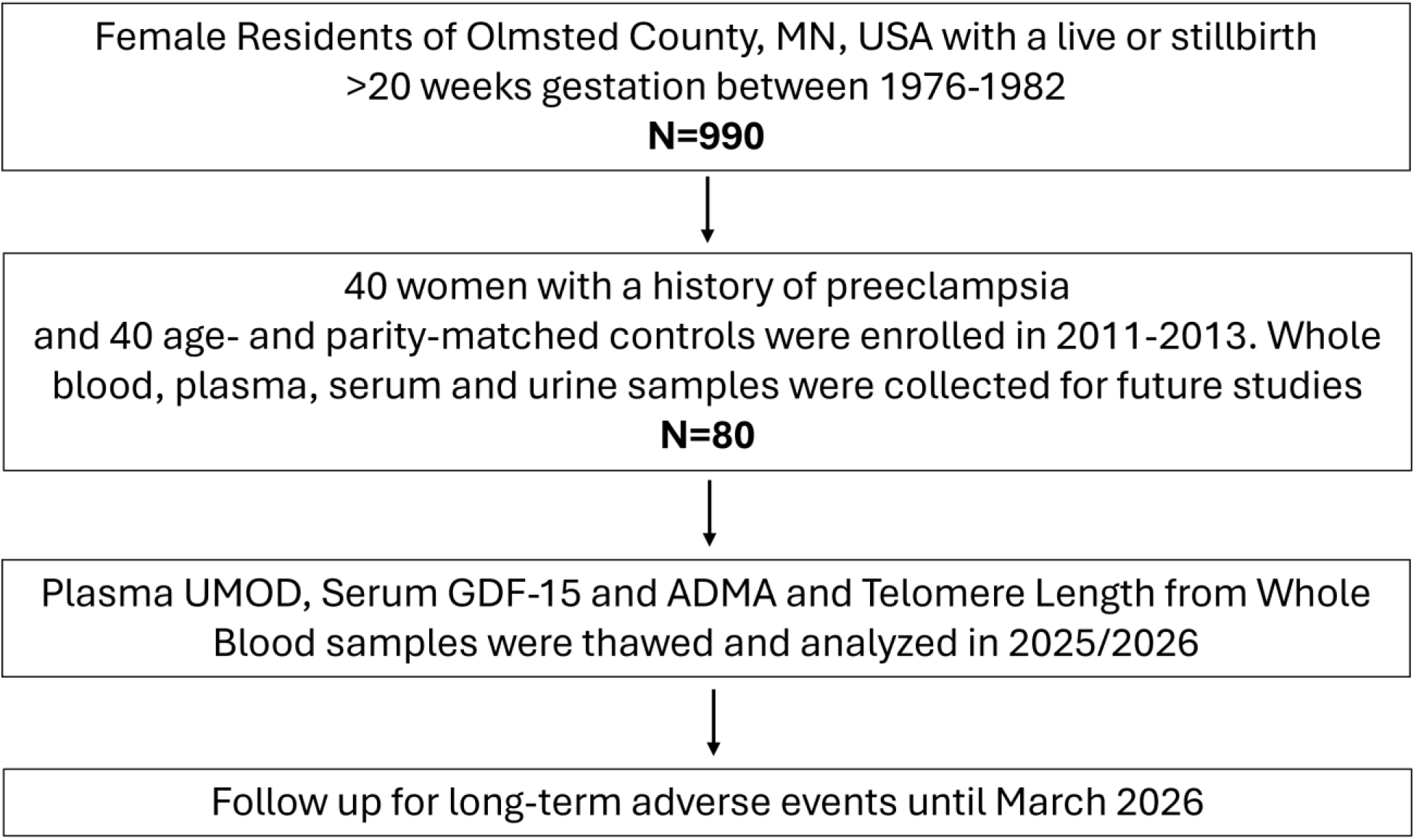
Flowchart of inclusion and exclusion criteria and Experiments Timeline All 80 women enrolled in the study had telomere lengths available for analysis. Due to inadequate voids, n=75 (37 PE and 38 controls) had plasma uromodulin and serum GDF-15 measurements available, and n=76 (38 PE and 38 controls) had ADMA serum levels available.

All participants provided written informed consent at enrollment between 1/1/2011 and 12/31/2013, at which time blood samples were collected.

### Blood collection and preparation for this study

Blood samples were obtained following an overnight fast, aliquoted and collected in 3-4 EDTA-containing tubes. First, the whole blood was frozen at -80°C. After centrifugation, plasma and serum were separated from additional blood samples and stored at -80°C until thawed. To ensure blind analyses, all clinical identifiers were removed before the experiments were performed.

### Plasma/Serum GDF15, UMOD and ADMA Analysis

Plasma UMOD concentrations were measured using a commercially available enzyme-linked immunosorbent assay (ELISA) kit Bio Vendor (catalog No. RD191163200R Brno, Czech Republic), while serum levels of GDF-15 and ADMA were measured by Abcam (catalog No. ab155432, Cambridge, UK) and Invitrogen (catalog No. EEL009, Carlsbad, CA, USA) ELISA Kits, respectively.

### Telomere Length Measurement

DNA from whole blood was extracted using the DNeasy Blood & Tissue Kit (QIAGEN) following the manufacturer’s protocol, quantified by a NanoDrop spectrophotometer (Thermo Fisher Scientific) and stored at −20 °C. Telomere lengths were evaluated by qPCR using Absolute Human Telomere Length Quantification qPCR Assay Kits (ScienCell Research Laboratories, catalog No. 8918, Carlsbad, CA, USA) according to manufacturer instructions. In brief, for each DNA sample, two consecutive reactions were performed: the first to amplify a single-copy reference (SCR) gene and the second for the telomere sequence. The SCR primer set recognizes and amplifies a 100 bp-long region on human chromosome 17 and serves as a reference for data normalization. Both PCRs were performed in a final volume of 20 μL using 1 μL of reference/genomic DNA samples from cases and controls (5 ng), 2 μL of primer stock solution (telomere or SCR), 10 μL of Master Mix and 7 μL of nuclease-free water. The PCR conditions were as follows: 95°C for 10 minutes followed by 32 cycles of: 95°C for 20 seconds, 52°C for 20 seconds and 72°C for 45 seconds. All reactions were run in duplicate, and data were analyzed using qPCR software and the equation from the manufacturer’s manual as described elsewhere.^14,15^

### Assessment of coronary artery calcification (CAC)

CAC images were obtained using a 64 detector computed tomography scanner (Siemens Sensation 64, Siemens Medical Solutions, Forcheim, Germany), with a scan configured to cover the heart as described previously, while calcium scoring was performed using the Agatston scoring method as described previously.^11,12^

### Statistical Analysis

Continuous variables are presented as mean (standard deviation [SD]), and categorical variables are summarized as counts and percentages. Because GDF-15 and ADMA measurements exhibited non-normal distributions, values were transformed for all subsequent analyses using a log base 2 transformation. Association analysis between PE status and plasma UMOD, GDF-15, ADMA and telomere length was performed both unadjusted using the equal variance t-test, as well as using analysis of covariance (ANCOVA) models adjusted for BMI and use of anti-hypertensive medications, to assesses whether PE-biomarker associations persist after accounting for important late-life cardiometabolic characteristics that may also lie on the causal pathway. Additional analyses looking at subtypes of PE and normotensive women with and without hypertension later in life, were also conducted using ANOVA models for unadjusted comparisons and ANCOVA models as described previously for adjusted comparisons. Correlations between these measures and predictors/markers of increased cardiovascular risk (coronary artery calcification [CAC], atherosclerosis and chronic arterial inflammation [high-sensitivity C-reactive protein (hs-CRP)] and insulin resistance (Homeostatic Model Assessment of Insulin Resistance [HOMA-IR]), were assessed using Spearman’s rank correlation coefficients.

The Kaplan-Meier method and the univariable Cox proportional hazards model were used to study the association between PE status and subsequent incidence of the composite outcome (CAD, HF, TIA, stroke, or CKD). Two patients with CKD at the time of enrollment were excluded from the analysis to evaluate *de novo* conditions. In addition, the association between plasma UMOD, GDF-15, ADMA, and telomere length and the incidence of the composite event was also evaluated using both unadjusted and adjusted Cox proportional hazards models; adjustments were made using the same variables used to evaluate associations across PE status. In all analyses, patients were followed from the enrollment date to the date of the condition diagnosis or were censored at the time of the last clinical visit recorded in the Rochester Epidemiology Project or death, whichever came first. Models were fit using age as the time scale, with women entering the risk set at their ages at study enrollment.

Results were considered statistically significant at the 0.05 alpha level. All analyses were performed using R version 4.4.1 (R Foundation for Statistical Computing, Vienna, Austria).

## RESULTS

### Plasma UMOD, GDF-15, ADMA, and telomere length in PE vs control subjects

Demographic and clinical characteristics of study subjects at baseline are shown in **Supplementary Table S1**. Patients with a PE history had significantly increased BMIs (median [IQR] 29.8 (25.9, 33.7) vs 25.3 (23,1, 32.0); p=0.023), hs-CRP ((median (IQR) 0.10 (0.05, 0.176) vs 0.18 (0.11, 0.32.); p=0.002), HOMA-IR ((median (IQR) 1.07 (0.76, 1.47) vs 1.829 (1.05, 3.98)) and a significantly higher percentage used anti-hypertensive therapy (n=23, 58%), compared to patients with a history of normotensive pregnancies (n=5, 13%); p<0.001), while lipid profiles, Framingham Risk Score, serum creatinine and eGFR were not significantly different between the groups at the time of enrollment as reported in previous studies performed on this cohort.^10,12^

All 80 women enrolled in the study had telomere lengths available for analysis. Due to inadequate voids, n=75 (37 PE and 38 controls) had plasma UMOD and serum GDF-15 measurements available, and n=76 (38 PE and 38 controls) had ADMA serum levels available. Women with a PE history had significantly lower levels of plasma UMOD compared to normotensive women (mean (SD): 124 (62) vs 167 (59), respectively, P=0.003) (see **Table 1** and **Figure 2**); telomere lengths were also found to be significantly reduced in the PE group compared to the normotensive group (mean (SD) 6.62 (1.68) vs 7.67 (1.62), respectively, P=0.006). Conversely, GDF-15 and ADMA levels were both significantly elevated in women with PE compared to controls (mean (SD) Log2(GDF-15): 9.68 (0.87) vs 9.33 (0.63), P=0.045 and mean (SD) Log2(ADMA): 6.83 (0.52) vs 6.49 (0.50), P=0.005). All results except GDF-15 remained statistically significant after adjustments for BMI and anti-hypertensive medication use (see **Table 1**). See **Supplementary Table S2** for comparisons across both PE and hypertension subtypes.

**Figure 2.**
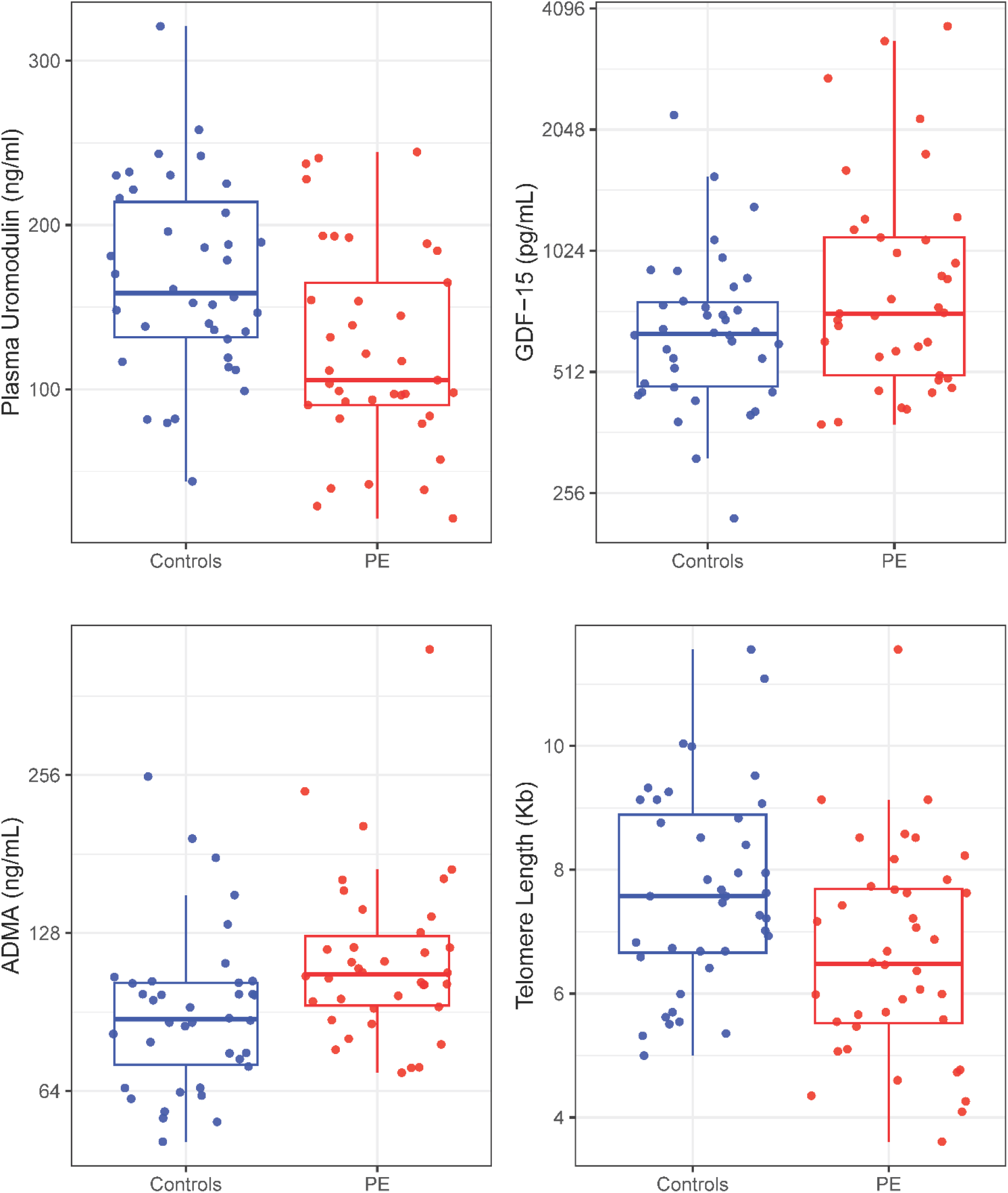
Plasma UMOD, GDF-15, ADMA, and Telomere length levels in women with and without a history of PE GDF-15 and ADMA levels were presented on the log base 2 scale for the y-axis.

**Table 1.** Unadjusted and Adjusted Associations between plasma UMOD, GDF15, ADMA and telomere length and PE status

|  | **Controls** |  | **PE** |  | **Unadjusted P value** | **Adjusted P value** |
|---|---|---|---|---|---|---|
| **UMOD Measure** | **N** | **Mean (SD)** | **N** | **Mean (SD)** |  |  |
| Plasma UMOD (ng/mL) | 38 | 167 (59) | 37 | 124 (62) | **0.003** | **0.041** |
| Log<sub>2</sub>(GDF15 (pg/mL)) | 38 | 9.33 (0.63) | 37 | 9.68 (0.87) | **0.045** | 0.37 |
| Log<sub>2</sub>(ADMA (ng/mL)) | 38 | 6.49 (0.50) | 38 | 6.83 (0.52) | **0.005** | **0.009** |
| Telomere length (Kb) | 40 | 7.67 (1.62) | 40 | 6.62 (1.68) | **0.006** | **0.033** |

Unadjusted P value = derived from equal variance T-test.
Adjusted P-value = Derived from ANCOVA model adjusted for the covariates BMI and anti-hypertensive medication.
P-values in bold denote statistical significance at the 0.05 alpha level.

### Correlations between Plasma UMOD, GDF-15, ADMA, and telomere length and predictors of increased CVD risk, vascular inflammation and atherosclerosis and insulin resistance

When assessing correlations between plasma UMOD, GDF-15, ADMA, and telomere length, only GDF-15 and ADMA exhibited a significant, positive correlation with each other (Spearman’s rho: 0.26 (95% CI, 0.02, 0.47), P=0.027). For plasma uromodulin, a modest negative correlation was observed with GDF-15 (r:-0.22 (95% CI -0.43, 0.02)) and a modest positive correlation with telomere length (r: 0.22 (95% CI -0.01, 0.43)), although results approached, but did not meet, the traditional threshold for statistical significance (see **Supplemental Table S3).** In **Table 2**, Spearman correlation coefficient estimates plasma UMOD, GDF-15, ADMA, and telomere lengths compared to CAC, hs-CRP, and HOMA-IR. GDF-15 was positively correlated with CAC (ρ=0.39) and HOMA-IR (ρ=0.25), whereas ADMA showed weaker positive correlations with CAC (ρ=0.23) and HOMA-IR (ρ=0.24). UMOD was inversely correlated with CAC (ρ=−0.25), hs-CRP (ρ=−0.29), and HOMA-IR (ρ=−0.40). Telomere length was inversely correlated with HOMA-IR (ρ=−0.32).

**Table 2.** Correlations between Plasma UMOD, GDF-15, ADMA, and telomere lengths and predictors of increased CVD risk (CAC), vascular inflammation and atherosclerosis (hs-CRP) and insulin resistance (HOMA-IR)

| Variable 1 | Variable 2 | N | Spearman’s rho (95% CI) | P |
|---|---|---|---|---|
| Log<sub>2</sub>(GDF-15 (pg/mL)) | CAC | 74 | 0.39 (0.17, 0.57) | **<0.001** |
| Log<sub>2</sub>(GDF-15 (pg/mL)) | hs-CRP | 69 | 0.11 (-0.14, 0.34) | 0.38 |
| Log<sub>2</sub>(GDF-15 (pg/mL)) | HOMA-IR | 75 | 0.25 (0.02, 0.46) | **0.031** |
| Log<sub>2</sub>(ADMA (ng/mL)) | CAC | 75 | 0.23 (0.00, 0.44) | **0.043** |
| Log<sub>2</sub>(ADMA (ng/mL)) | hs-CRP | 71 | 0.02 (-0.22, 0.26) | 0.87 |
| Log<sub>2</sub>(ADMA (ng/mL)) | HOMA-IR | 76 | 0.24 (0.01, 0.45) | **0.033** |
| Plasma Uromodulin (ng/ml) | CAC | 74 | -0.25 (-0.46, -0.01) | **0.032** |
| Plasma Uromodulin (ng/ml) | hs-CRP SCOR | 69 | -0.29 (-0.50, -0.05) | **0.014** |
| Plasma Uromodulin (ng/ml) | HOMA-IR | 75 | -0.40 (-0.58, -0.18) | **<0.001** |
| Telomere Length (Kb) | CAC | 79 | -0.14 (-0.36, 0.09) | 0.21 |
| Telomere Length (Kb) | hs-CRP SCOR | 74 | -0.21 (-0.42, 0.03) | 0.073 |
| Telomere Length (Kb) | HOMA-IR | 80 | -0.32 (-0.51, -0.10) | **0.004** |

Spearman’s rank correlation coefficients were reported for all analyses.
P-values in bold denote statistical significance at the 0.05 alpha level.
CAC=Coronary artery calcification; hs-CRP=high-sensitivity C-reactive protein; HOMA-IR=Homeostatic Model Assessment of Insulin Resistance.

### PE vs control subjects and Risk of CVD and CKD through prolonged follow-up

Among the 78 women without prevalent CKD at enrollment (2 women with CKD were excluded to account for known CVD risks associated with kidney disease), there was a total of 32 incident composite events (15 with CAD, 3 with HF, 3 with TIA/stroke, and 11 with CKD) over a median (95% CI) follow-up of 11.2 (11.1, 11.4) years. Women with a PE history demonstrated an increased risk of developing one of the events (CAD, HF, TIA, stroke, or CKD), compared to normotensive women (hazard ratio, 2.43 [95% CI, 1.17-5.06]; P=0.017; see **Figure 3**) in this extended timeframe.

**Figure 3.**
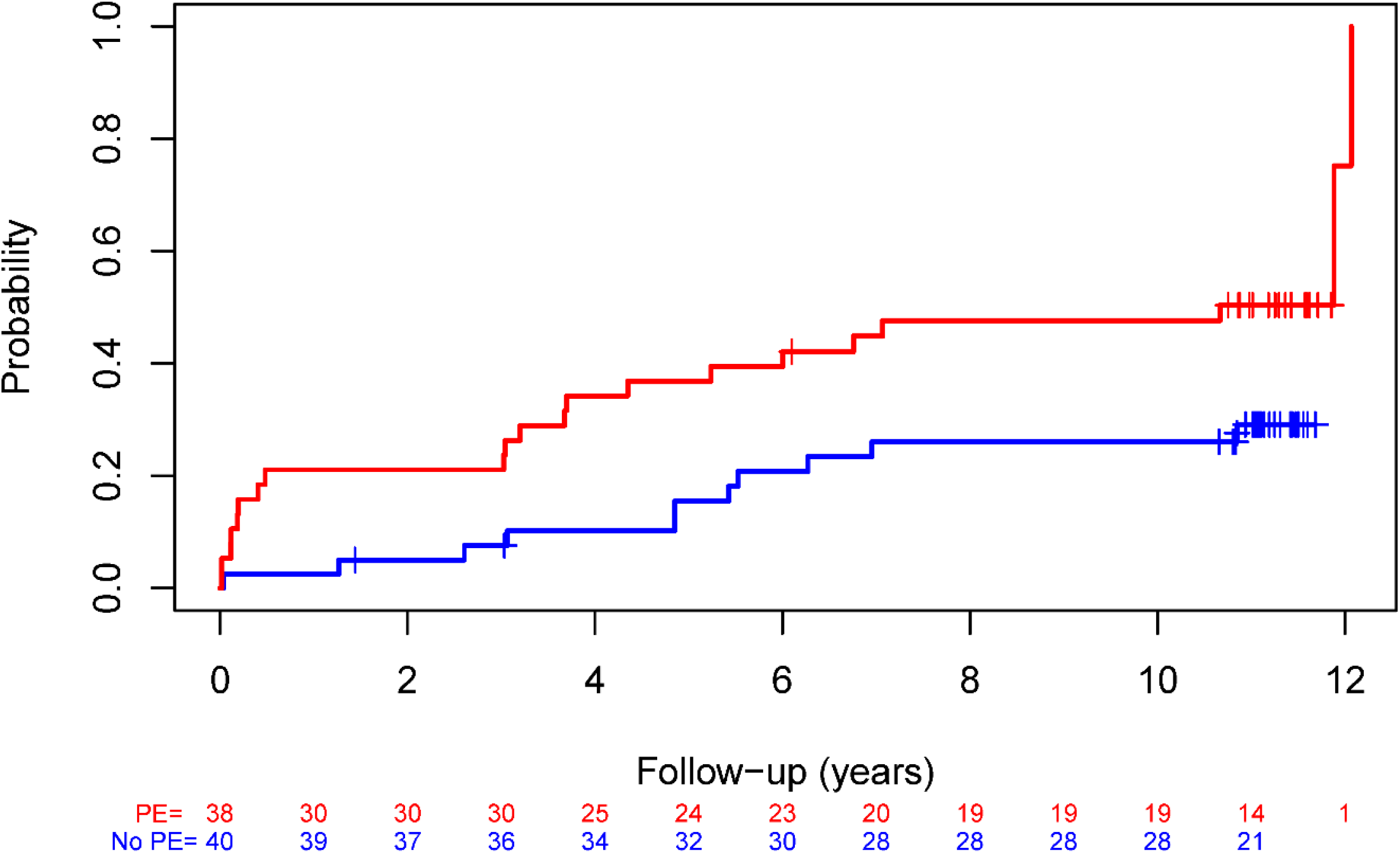
Cumulative incidence of time to the composite event of coronary artery disease (CAD), heart failure (HF), transient ischemic attack (TIA)/ stroke, or chronic kidney disease (CKD), by PE status over an extended follow-up up to March 2026

### Plasma UMOD, GDF-15, ADMA, and telomere length and Risk of future CVD and CKD

In an unadjusted analysis, women with higher plasma UMOD levels demonstrated a significantly lower risk of developing one of the events (CAD, HF, TIA, stroke, or CKD) (Hazard Ratio, 0.94 [95% CI, 0.89–0.999]; P=0.045); the inverse association between UMOD and the composite outcome was attenuated after adjustment. Increased GDF-15 levels were found to be significantly associated with a higher risk of this composite event (Hazard Ratio, 2.04 [95% CI, 1.34-3.10]; P<0.001), and results were similar after adjusting for BMI and anti-hypertensive medication use (Hazard Ratio, 2.08 [95% CI, 1.31-3.28]; P=0.002). Associations of ADMA and telomere length with the composite outcome did not meet the prespecified threshold for statistical significance (see **Table 3**).

**Table 3.** Cox regression models looking at time to CVD/CKD vs plasma UMOD, ADMA, GDF-15, and telomere length

|  |  |  | **Unadjusted** |  | **Adjusted** |  |
|---|---|---|---|---|---|---|
| **Variable** | **N** | **E** | **HR (95% CI)** | **P-value** | **HR (95% CI)** | **P-value** |
| Plasma Uromodulin (per 10 ng/ml) | 73 | 31 | 0.94 (0.89-0.999) | **0.045** | 0.98 (0.91, 1.05) | 0.59 |
| Log<sub>2</sub>(GDF-15 (pg/mL)) | 73 | 30 | 2.04 (1.34–3.10) | **<0.001** | 2.08 (1.31, 3.28) | **0.002** |
| Log<sub>2</sub>(ADMA (ng/mL)) | 74 | 31 | 1.64 (0.88–3.06) | 0.12 | 1.58 (0.83, 3.01) | 0.17 |
| Telomere length | 78 | 32 | 0.89 (0.71-1.12) | 0.32 | 0.97 (0.76-1.23) | 0.79 |

Adjusted models included adjustments for the covariates BMI and anti-hypertensive medication. P-values in bold denote statistical significance at the 0.05 alpha level.

## DISCUSSION

Preeclampsia is a heterogeneous disorder involving multiple pathophysiologic pathways, but whether abnormalities representing these pathways persist decades after the affected pregnancy and contribute similarly to long-term cardiorenal risk is unknown. In this study of postmenopausal women evaluated approximately 3 decades after pregnancies complicated by PE or normotensive pregnancies, we found evidence of persistent differences across several biological domains. Women with prior PE had lower circulating UMOD, higher ADMA, and shorter leukocyte telomere length after adjustments for BMI and anti-hypertensive medication use, consistent with long-term differences in pathways related to renal tubular integrity, nitric oxide metabolism, and biological aging. In contrast, the difference in GDF-15, a marker of cellular stress, was attenuated after adjustment. During a median of 11.2 years of subsequent follow-up, prior PE was associated with a 2.43-fold higher hazard of incident cardiovascular or kidney disease. Importantly, the relationships between individual biomarkers and subsequent outcomes differed: GDF-15 demonstrated the strongest association with incident clinical events, whereas the other biomarkers primarily distinguished women according to prior PE status. These findings suggest that biological pathways implicated in PE may differ both in their persistence after pregnancy and in their relationships with long-term cardiorenal outcomes.

Renal tubular dysfunction represents one pathway implicated in PE. UMOD is produced exclusively by epithelial cells of the thick ascending limb and distal nephron, with small amounts being released into the circulation through the basolateral membrane. Both urinary and circulating UMOD reflect functional tubular mass and renal parenchymal integrity.^3,11,16^ Emerging evidence indicates plasma UMOD as a more robust, consistent, and reliable biomarker of kidney diseases and CVD. Urinary UMOD, although much more abundant, is subject to large fluctuations and enzyme degradation and requires very careful processing and sample storage.^17^ We therefore measured plasma UMOD to examine whether a renal tubular phenotype associated with PE remains detectable decades after pregnancy. A higher level of UMOD is representative of healthy, functional kidney tubular mass and is associated with better kidney function.^18^ In this study, as we predicted, women with a PE history had significantly lower levels of plasma UMOD. Moreover, UMOD levels negatively correlated with CAC, hs-CRP and HOMA-IR, consistent with prior observations linking higher circulating UMOD with more favorable renal and cardiovascular phenotypes.^16^ Women with a PE history with lower plasma UMOD levels also demonstrated a significantly increased risk of developing one of the CVD or CKD events, although this association was attenuated after adjustments for BMI and anti-hypertensive medications. The attenuation of results after adjustment could be for multiple reasons. One is the small sample size of this study which underpowered the results. It is also possible that BMI and hypertension are mediators for the effect of PE on UMOD concentrations. A history of preeclampsia has been associated with a greater future BMI and chronic endothelial/vascular stress and hypertension, which can drive chronic kidney tubule stress and reduce UMOD levels.^19–21^ The attenuation after adjustment suggests that the association between UMOD and clinical outcomes may be explained partly by late-life cardiometabolic characteristics, although limited statistical power precludes firm conclusions.

GDF-15, a distant member of the transforming growth factor-β (TGF-β) cytokine superfamily, has widely been established as a biomarker for the severity of multiple disease conditions. It is heavily involved in stress signaling and its circulating levels rise in response to metabolic stress and obesity.^22^ In the cardiovascular system, GDF-15 is secreted locally by cardiac myocytes and vascular cells in response to pressure overload, high oxidative stress and ischemia to protect the tissues with its anti-proliferative, anti-inflammatory and anti-apoptotic actions.^23,24^ Elevated systemic levels of GDF-15 are a strong predictor of CVDs and accelerated aging, with GDF-15 being one of the key components of the senescence-associated secretory phenotype (SASP).^23,25^ GDF-15 is linked to morning sickness during pregnancy and higher levels are observed preceding a diagnosis of PE, where it may serve as a clinical biomarker for early PE detection.^4,26,27^ The present study revealed significantly increased levels of GDF-15 in women with a PE history. Moreover, elevated levels of GDF-15 were strongly associated with adverse clinical outcomes and were positively correlated with CAC and HOMA-IR. Altogether, these findings indicate alterations in cellular stress response especially in the cardiovascular system, impaired response to metabolic stress, and accelerated aging in postmenopausal women with a history of PE.

Endothelial dysfunction and impaired nitric oxide (NO) bioavailability are central features of PE. ADMA, an endogenous inhibitor of nitric oxide synthase, is elevated during pregnancies complicated by PE and provides a circulating marker of altered NO biology.^5,28,29^ In the present study, ADMA remained higher decades after pregnancy among women with prior PE, even after adjustment for BMI and anti-hypertensive medication use. Its modest correlations with CAC and HOMA-IR further support an association with late-life vascular and metabolic phenotypes. However, ADMA was not significantly associated with subsequent composite cardiorenal events. Thus, persistent abnormalities in NO-related biology may characterize women with prior PE without necessarily serving as a strong late-life prognostic marker in this cohort.

Biological aging represents another potential pathway linking PE with later-life disease. Shorter telomere length has been reported in association with PE,^6^ and leukocyte telomere length provides a marker of cumulative replicative and cellular stress.^30,31^ Women with prior PE in our study had shorter leukocyte telomeres approximately 3 decades after pregnancy, consistent with persistent differences in biological aging. Because telomere length was measured at a single time point, however, our findings cannot establish accelerated telomere attrition. Telomere length was associated with HOMA-IR but not with subsequent composite cardiorenal events, suggesting that this late-life aging phenotype may characterize prior PE without independently identifying near-term clinical risk in this small cohort.

In conclusion, in postmenopausal women evaluated decades after pregnancy, a history of PE was associated with persistent differences in biomarkers representing several biological pathways implicated in the disorder, including renal tubular integrity, nitric oxide metabolism, and biological aging. Importantly, these pathway-specific biomarkers showed different relationships with subsequent disease: UMOD, ADMA, and telomere length primarily distinguished women with prior PE from those with normotensive pregnancies, whereas GDF-15, a marker of cellular stress, showed the strongest association with subsequent cardiorenal events. These findings support the concept that the biological heterogeneity of PE may extend beyond pregnancy and that different pathways associated with the disorder may have distinct relationships with long-term cardiovascular and kidney risk. Larger longitudinal studies are needed to determine whether pathway-specific biomarkers can improve risk stratification after PE and to define the mechanisms linking heterogeneous PE phenotypes with later-life disease.

### Study limitations

Our study has some important limitations which should be considered while interpreting the results. First, the study population was small, which resulted in limited statistical power for complex, large multivariable analyses. Second, this study was conducted at a single center reflecting the population of Olmsted County, Minnesota, which was primarily white and non-Hispanic at the time when study participants gave birth; therefore, these results may not be generalizable to women of other ethnicities and races, especially as the risks of PE and hypertension have been demonstrated to vary with ethnicity.^32^ Third, we measured plasma/serum GDF-15, ADMA and UMOD in samples that were collected and stored for approximately 15 years before the analysis. However, previous studies have suggested that circulating UMOD, GDF15 and ADMA are highly stable antigens with a high concordance under various temperatures even after five freeze and thaw cycles.^33–36^ In the present study, available aliquots used for analysis were thawed for the first or second time. Fourth, all women with severe cardiovascular and renal events at the time of enrollment were excluded from the study. All participants were of relatively good health at the time of enrollment, which could underestimate the overall influence of PE on kidney injury and cardiovascular risk.

## Abbreviations used in text

PE: Preeclampsia
CVD: cardiovascular disease
CKD: chronic kidney disease
UMOD: uromodulin
GDF-15: growth differentiation factor-15
ADMA: asymmetric dimethylarginine
BMI: body mass index
eGFR: estimated glomerular filtration rate
CAC: coronary artery calcification
hs-CRP: high-sensitivity C-reactive protein
HOMA-IR: homeostatic Model Assessment of Insulin Resistance

---

**Supplementary Table S1.**
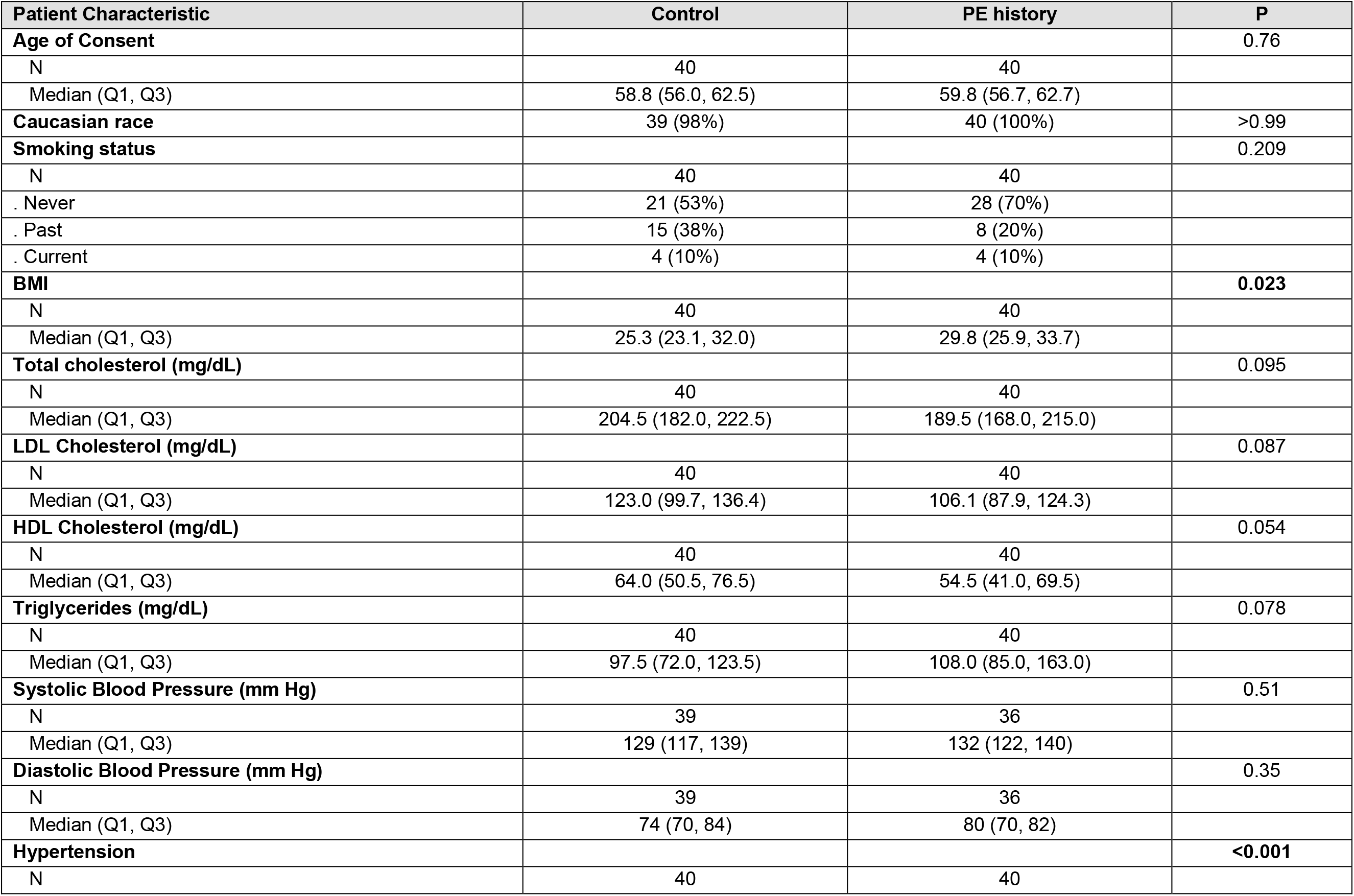
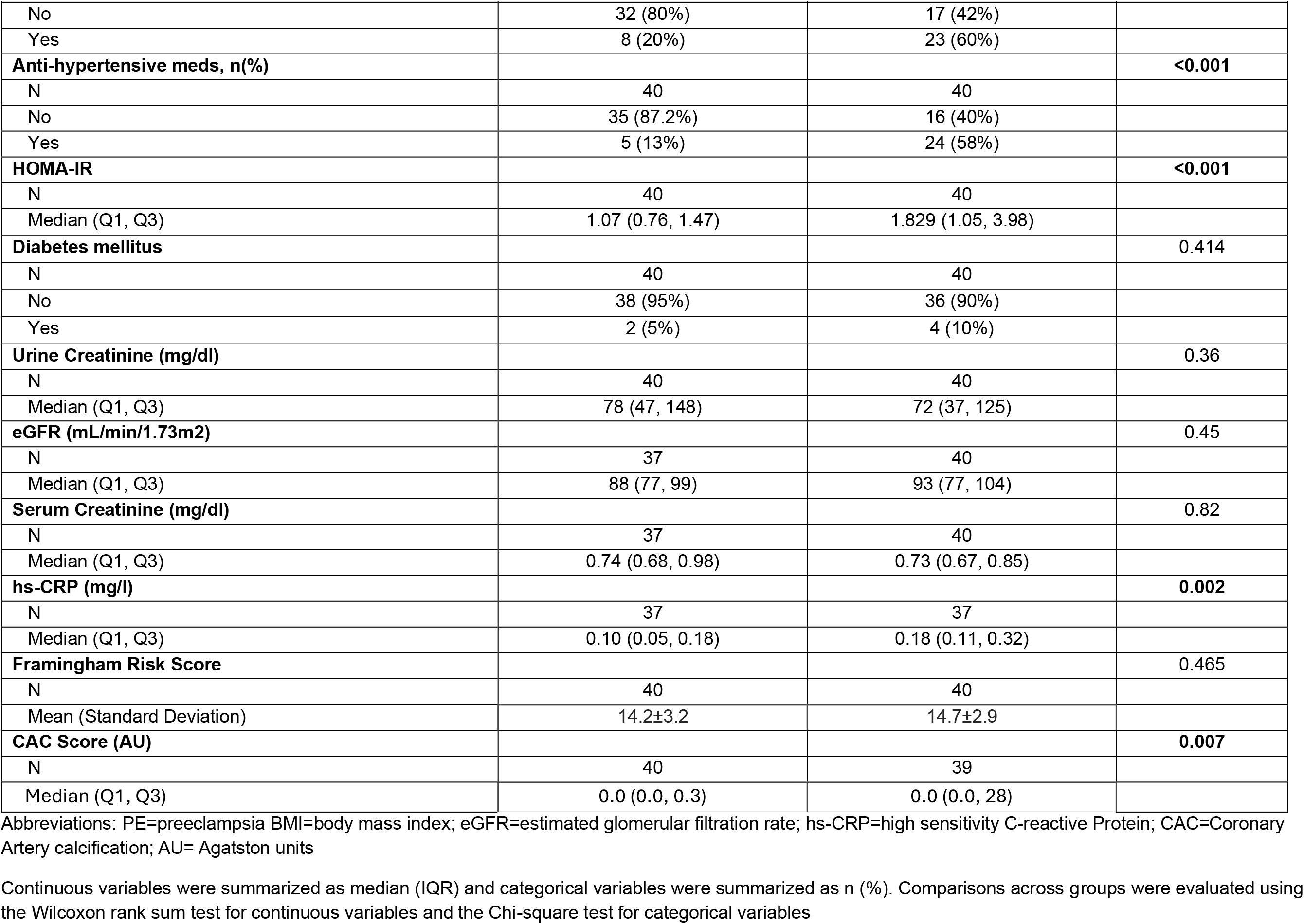
Demographics and Clinical Characteristics of Study Participants

| Patient Characteristic | Control | PE history | P |
|---|---|---|---|
| **Age of Consent** |  |  | 0.76 |
| N | 40 | 40 |  |
| Median (Q1, Q3) | 58.8 (56.0, 62.5) | 59.8 (56.7, 62.7) |  |
| **Caucasian race** | 39 (98%) | 40 (100%) | >0.99 |
| **Smoking status** |  |  | 0.209 |
| N | 40 | 40 |  |
| . Never | 21 (53%) | 28 (70%) |  |
| . Past | 15 (38%) | 8 (20%) |  |
| . Current | 4 (10%) | 4 (10%) |  |
| **BMI** |  |  | **0.023** |
| N | 40 | 40 |  |
| Median (Q1, Q3) | 25.3 (23.1, 32.0) | 29.8 (25.9, 33.7) |  |
| **Total cholesterol (mg/dL)** |  |  | 0.095 |
| N | 40 | 40 |  |
| Median (Q1, Q3) | 204.5 (182.0, 222.5) | 189.5 (168.0, 215.0) |  |
| **LDL Cholesterol (mg/dL)** |  |  | 0.087 |
| N | 40 | 40 |  |
| Median (Q1, Q3) | 123.0 (99.7, 136.4) | 106.1 (87.9, 124.3) |  |
| **HDL Cholesterol (mg/dL)** |  |  | 0.054 |
| N | 40 | 40 |  |
| Median (Q1, Q3) | 64.0 (50.5, 76.5) | 54.5 (41.0, 69.5) |  |
| **Triglycerides (mg/dL)** |  |  | 0.078 |
| N | 40 | 40 |  |
| Median (Q1, Q3) | 97.5 (72.0, 123.5) | 108.0 (85.0, 163.0) |  |
| **Systolic Blood Pressure (mm Hg)** |  |  | 0.51 |
| N | 39 | 36 |  |
| Median (Q1, Q3) | 129 (117, 139) | 132 (122, 140) |  |
| **Diastolic Blood Pressure (mm Hg)** |  |  | 0.35 |
| N | 39 | 36 |  |
| Median (Q1, Q3) | 74 (70, 84) | 80 (70, 82) |  |
| **Hypertension** |  |  | **<0.001** |
| N | 40 | 40 |  |
| No | 32 (80%) | 17 (42%) |  |
| Yes | 8 (20%) | 23 (60%) |  |
| **Anti-hypertensive meds, n(%)** |  |  | **<0.001** |
| N | 40 | 40 |  |
| No | 35 (87.2%) | 16 (40%) |  |
| Yes | 5 (13%) | 24 (58%) |  |
| **HOMA-IR** |  |  | **<0.001** |
| N | 40 | 40 |  |
| Median (Q1, Q3) | 1.07 (0.76, 1.47) | 1.829 (1.05, 3.98) |  |
| **Diabetes mellitus** |  |  | 0.414 |
| N | 40 | 40 |  |
| No | 38 (95%) | 36 (90%) |  |
| Yes | 2 (5%) | 4 (10%) |  |
| **Urine Creatinine (mg/dl)** |  |  | 0.36 |
| N | 40 | 40 |  |
| Median (Q1, Q3) | 78 (47, 148) | 72 (37, 125) |  |
| **eGFR (mL/min/1.73m2)** |  |  | 0.45 |
| N | 37 | 40 |  |
| Median (Q1, Q3) | 88 (77, 99) | 93 (77, 104) |  |
| **Serum Creatinine (mg/dl)** |  |  | 0.82 |
| N | 37 | 40 |  |
| Median (Q1, Q3) | 0.74 (0.68, 0.98) | 0.73 (0.67, 0.85) |  |
| **hs-CRP (mg/l)** |  |  | **0.002** |
| N | 37 | 37 |  |
| Median (Q1, Q3) | 0.10 (0.05, 0.18) | 0.18 (0.11, 0.32) |  |
| **Framingham Risk Score** |  |  | 0.465 |
| N | 40 | 40 |  |
| Mean (Standard Deviation) | 14.2±3.2 | 14.7±2.9 |  |
| **CAC Score (AU)** |  |  | **0.007** |
| N | 40 | 39 |  |
| Median (Q1, Q3) | 0.0 (0.0, 0.3) | 0.0 (0.0, 28) |  |

Abbreviations: PE=preeclampsia BMI=body mass index; eGFR=estimated glomerular filtration rate; hs-CRP=high sensitivity C-reactive Protein; CAC=Coronary Artery calcification; AU= Agatston units
Continuous variables were summarized as median (IQR) and categorical variables were summarized as n (%). Comparisons across groups were evaluated using the Wilcoxon rank sum test for continuous variables and the Chi-square test for categorical variables

**Supplemental Table S2.**
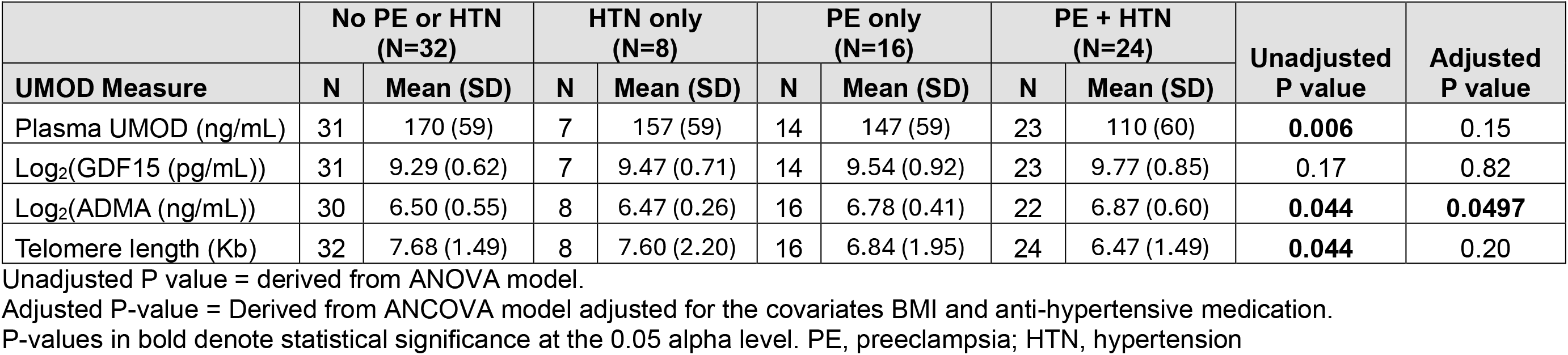
Unadjusted and Adjusted Associations between plasma UMOD, GDF15, ADMA and telomere length and PE and hypertension status

**Supplemental Table S3.** Correlations between plasma UMOD, GDF15, ADMA and telomere length

| **Variable 1** | **Variable 2** | **N** | **Spearman’s rho (95% CI)** | **P** |
|---|---|---|---|---|
| Log<sub>2</sub>(GDF-15 (pg/mL) | Log<sub>2</sub>(ADMA (ng/mL)) | 71 | 0.26 (0.02, 0.47) | **0.027** |
| Log<sub>2</sub>(GDF-15 (pg/mL)) | Plasma Uromodulin (ng/ml) | 71 | -0.22 (-0.43, 0.02) | 0.068 |
| Log<sub>2</sub>(GDF-15 (pg/mL)) | Telomere Length (Kb) | 75 | -0.06 (-0.29, 0.17) | 0.59 |
| Log<sub>2</sub>(ADMA (ng/mL)) | Plasma Uromodulin (ng/ml) | 71 | -0.10 (-0.33, 0.14) | 0.40 |
| Log<sub>2</sub>(ADMA (ng/mL)) | Telomere Length (Kb) | 76 | -0.08 (-0.30, 0.16) | 0.51 |
| Plasma Uromodulin (ng/ml) | Telomere Length (Kb) | 75 | 0.22 (-0.01, 0.43) | 0.053 |

Spearman’s rank correlation coefficients were reported for all analyses.
P-values in bold denote statistical significance at the 0.05 alpha level.

